# The interaction between NLRP1 and oxidized TRX1 involves a transient disulfide bond

**DOI:** 10.1101/2023.09.27.559829

**Authors:** Michael B. Geeson, Jeffrey C. Hsiao, Lydia P. Tsamouri, Daniel P. Ball, Daniel A. Bachovchin

## Abstract

NLRP1 is an innate immune receptor that detects pathogen-associated signals, assembles into a multiprotein structure called an inflammasome, and triggers a proinflammatory form of cell death called pyroptosis. We previously discovered that the oxidized, but not the reduced, form of thioredoxin-1 directly binds to NLRP1 and represses inflammasome formation. However, the molecular basis for NLRP1’s selective association with only the oxidized form of TRX1 has not yet been established. Here, we leveraged Alphafold-Multimer, site-directed mutagenesis, thiol-trapping experiments, and mass spectrometry to reveal that a specific cysteine residue (C427 in humans) on NLRP1 forms a transient disulfide bond with oxidized TRX1. Overall, this work demonstrates how NLRP1 monitors the cellular redox state, further illuminating an unexpected connection between the intracellular redox potential and the innate immune system.

## INTRODUCTION

NLRP1 was the first pattern-recognition receptor discovered that oligomerizes into an inflammasome ^1^. The human NLRP1 protein has an N-terminal pyrin domain (PYD) preceding nucleotide-binding (NACHT), leucine-rich repeat (LRR), function-to-find (FIIND), and caspase activation and recruitment (CARD) domains (**Fig. 1A**) ^2,3^. The FIIND undergoes post-translational autoproteolysis between its ZU5 and UPA subdomains, creating N-terminal (NT) and C-terminal (CT) fragments that remain associated and inactive ^4-6^. The proteasome-mediated degradation of the NT fragment releases the CT fragment from autoinhibition ^7,8^, but each freed CT fragment is then sequestered in a ternary complex with one full-length copy of NLRP1 and the serine peptidase DPP9 ^9,10^. Stimuli that accelerate NT degradation and/or destabilize the DPP9-mediated ternary complexes can liberate sufficient CT fragments to self-oligomerize and nucleate an inflammasome ^3,7,8,11^. However, the physiological-relevant danger signals that trigger these processes have still not been fully established.

**Figure 1.**
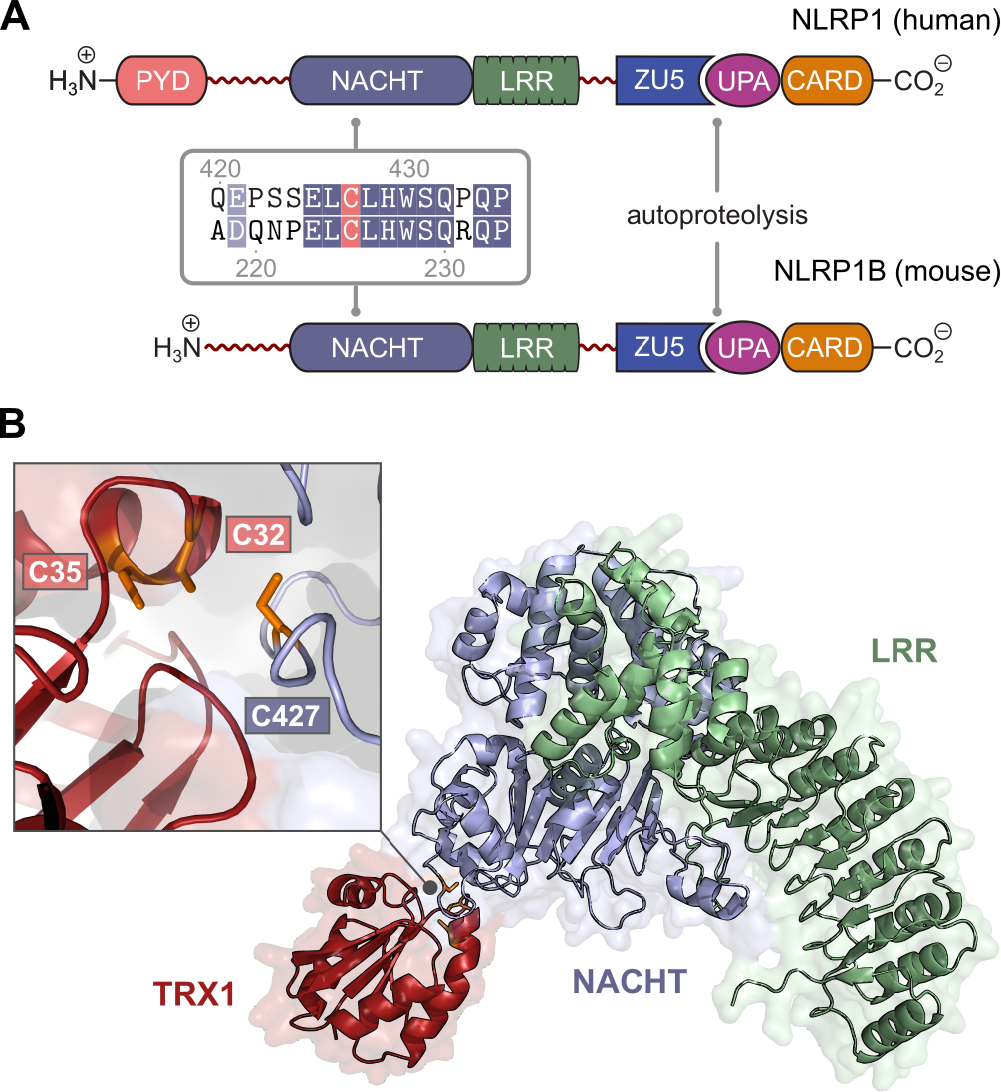
AlphaFold-Multimer predicts the protein-protein interface between NLRP1 and TRX1. (**A**) Domain architecture of human NLRP1 and mouse NLRP1B, indicating the sites of autoproteolysis, sequence alignment at the TRX1-binding interface, and the conserved cysteine residue. (**B**) Model of the NACHT-LRR-TRX1 heterodimer predicted by AlphaFold-Multimer. Inset: close-up view of the cysteine residues potentially involved in an intermolecular disulfide bond.

Two years ago, we reported that the oxidoreductase thioredoxin-1 (TRX1) directly binds to NLRP1 protein and suppresses its activation ^12,13^. TRX1, which plays a key role in the maintenance of the cytosolic redox environment, uses two active-site cysteines (C32 and C35) to catalyze the reduction of disulfide bonds ^14,15^. C32 in reduced TRX1 first reacts with an oxidized substrate to form an intermolecular disulfide bond. C35^TRX1^ then quickly attacks this disulfide bond, thereby reducing the target protein and oxidizing TRX1. We found that the oxidized (i.e., the disulfide), but not the reduced (i.e., the dithiol), form of TRX1 interacts with and stabilizes the folding of NLRP1’s NACHT-LRR region, thereby slowing NT degradation and repressing inflammasome formation. Overall, our results provided the seminal finding that a reduced cell state, which would eliminate the disulfide bond in oxidized TRX1, was critical for NLRP1 inflammasome activation.

Here, we wanted to determine the molecular basis for the selective association between NLRP1 and oxidized, but not reduced, TRX1. We first used the recently developed Alphafold-Multimer platform ^16^ to predict the structure of the NLRP1-TRX1 complex, which suggested that a hydrophobic ridge within the NLRP1 NACHT domain interacts with the active-site pocket of TRX1. Notably, this model predicted that C427 in human NLRP1 (C225 in mouse NLRP1B) is positioned close to C32 in TRX1. Using site-directed mutagenesis and thiol-trapping experiments, we discovered that C427 forms a transient disulfide bond with oxidized TRX1, essentially acting as a selectivity filter to bind only this form of the oxidoreductase. Overall, this work reveals a surprising molecular mechanism that links the intracellular redox state to the innate immune system.

## RESULTS

### Prediction of the TRX1-NLRP1 interaction

To explore the molecular basis of the interaction between TRX1 and the NACHT-LRR region (residues 229-990) of NLRP1, we first used AlphaFold-Multimer to predict the structure of the protein-protein complex ^16^. Although we were only able to use reduced TRX1 as an input due to the design of AlphaFold-Multimer, this platform nevertheless provided a model with a DockQ Score of 0.73, close to the cut-off for highly accurate scores (≥ 0.8) and well above the cut-off for incorrect scores (<0.23) (**Fig. 1B, Fig. S1**). Consistent with this high score, visual inspection of the predicted interface did not reveal any notable clashes between amino acid sidechains. The key binding interface involves a hydrophobic ridge on NLRP1 (residues ^426^LCLHW^430^) that fits into the active-site notch on TRX1 (residues ^31^WCGPC^35^). Intriguingly, this prediction placed C32 of TRX1 directly between C35 of TRX1 (2.7 Å) and C427 of NLRP1 (4.4 Å) (**Fig. 1B**, inset), raising the possibility that C427 of NLRP1 acts as a selectivity filter to restrict association to only the oxidized form of TRX1 by forming a transient disulfide bond. Beyond this potential intermolecular disulfide, salt bridges are predicted to form between R396^NLRP1^ and D60^TRX1^/D61^TRX1^ and E451^NLRP1^ and K36^TRX1^ (**Fig. S1A**).

We previously discovered that TRX1 also associates with rodent NLRP1 proteins, including mouse NLRP1B ^13^. We next similarly used AlphaFold-Multimer to predict the interaction between TRX1 and the NACHT-LRR (residues K30 to V810) of mouse NLRP1B (allele 1) ^17^. Gratifyingly, AlphaFold-Multimer returned a model with a DockQ score of 0.86, again indicative of a high-quality protein-protein interface. Moreover, the overlay of the human and the mouse interaction models resulted in excellent spatial alignment, particularly within TRX1 and the NACHT domain (**Fig. S1B**). Notably, C225 of mouse NLRP1 was predicted to interact in a similar way as C427 of human NLRP1 with the active site of TRX1.

### C427 mediates the NLRP1-TRX1 interaction

We next wanted to determine the importance of C427 in human NLRP1 and C225 in mouse NLRP1B for TRX1 binding. We therefore mutated these key cysteine residues to serine, along with several other cysteine residues on human NLRP1 as controls (C529, C754, and C757 were selected as controls because they are surface residues in the NACHT-LRR region). We then transiently expressed C-terminally FLAG-tagged wild-type and mutant proteins in HEK 293T cells, and enriched proteins from these lysates by anti-FLAG immunoprecipitation (IP) (**Fig. 2A**,**B**). Strikingly, we found that NLRP1^C427S^ did not bind TRX1, while the other three cysteine mutations in human NLRP1 retained binding (**Fig. 2A)**. Similarly, we found that mouse NLRP1B^C225S^ and the isolated human NACHT-LRR region (residues 229-990) with a C427S mutation did not bind TRX1 (**Fig. 2B**,**C**). As expected, DPP9, which associates with the NLRP1 FIIND domain ^9,10,18^, still bound to NLRP1^C427S^ and NLRP1B^C225S^. To further validate the AlphaFold-Multimer model and the importance of the NLRP1’s hydrophobic ridge for binding TRX1, we next mutated W430^NLRP1^, which occupies the central position in the interface (**Fig. S1A**), to glycine before performing a similar immunoprecipitation experiment. As expected, we found mutation of W430 completely abrogated the TRX1 interaction while retaining affinity for DPP9 (**Fig. S2**). Overall, these results show that C427^NLRP1^/C225 ^NLRP1B^ are required for the NLRP1-TRX1 interaction.

**Figure 2.**
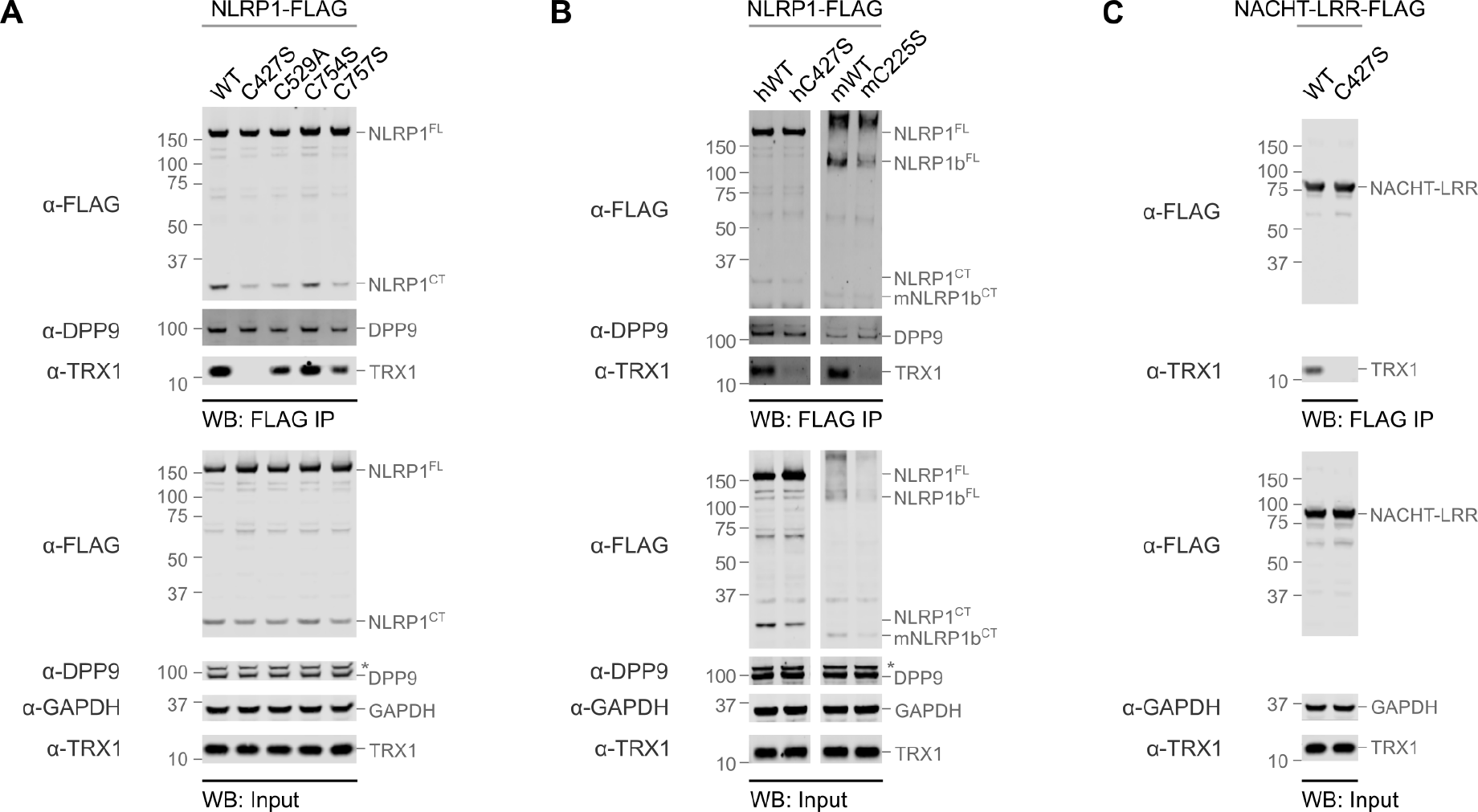
Mutation of C427 (human) or C225 (mouse) abrogates TRX1 binding to NLRP1. (**A-C**) HEK 293T cells were transiently transfected with plasmids encoding the indicated FLAG-tagged proteins before lysates were subjected to anti-FLAG IP and immunoblotting analyses.

### A transient intermolecular disulfide bond

As mentioned above, we hypothesized that C427^NLRP1^ might form a transient intermolecular disulfide bond with C32 on oxidized TRX1 that provides critical binding affinity. To explore this possibility, we next attempted to stabilize the transient NLRP1-TRX1 conjugate using thiol-trapping approaches (**Fig. 3A**). In these experiments, NLRP1-TRX1 complexes were treated with iodoacetamide (IAA) or N-ethyl maleimide (NEM), which react with free thiols. In theory, if C35^TRX1^ reacts with IAA or NEM while C32^TRX1^ is involved in a transient intermolecular bond with C427^NLRP1^, it will render the complex unable to resolve the disulfide between TRX1 and NLRP1 and thereby stabilize the interaction. Notably, such a disulfide-linked conjugate should be detectable by immunoblotting under non-reducing gel electrophoresis.

**Figure 3.**
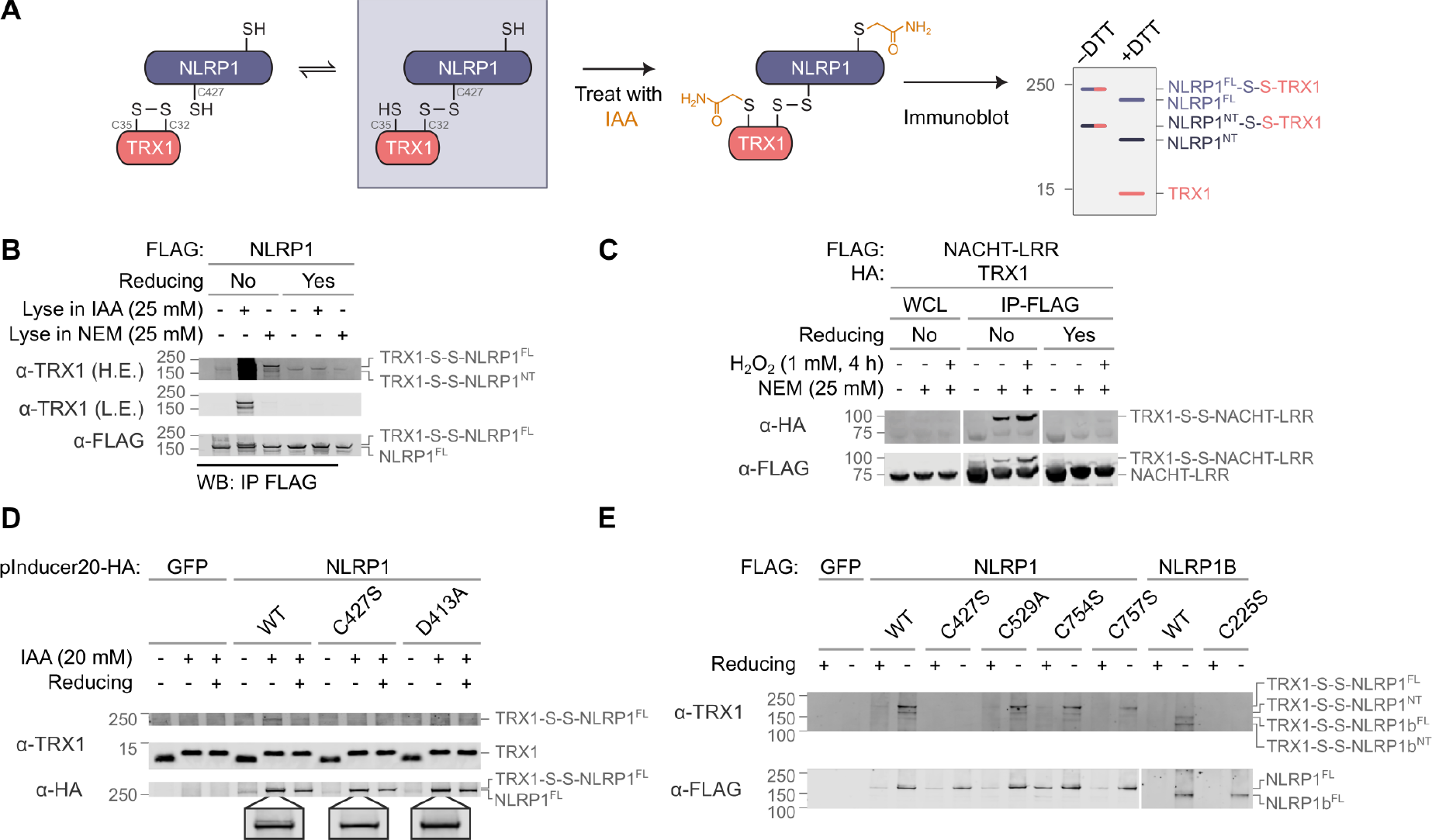
A transient intermolecular disulfide forms between NLRP1 and TRX1. **(A)** Schematic of the IAA- and NEM-conjugate trapping experiment. (**B**,**C**) HEK 293T cells were transiently transfected with plasmids encoding the indicated FLAG- or HA-tagged proteins before cells were lysed in IAA or NEM and proteins were enriched by anti-FLAG IP and analyzed by immunoblotting. In **C**, cells were treated with H_2_O_2_ as indicated prior to lysis. H.E., high exposure; L.E., low exposure. (**D**) *CARD8*^*-/-*^ THP-1 cells containing plasmids (pInducer-20 vector) encoding doxycycline-inducible, HA-tagged proteins were treated with DOX (1 ug/mL) for 16 h before being lysed in IAA and visualized by immunoblotting. (**E**) HEK 293T cells were transiently transfected with plasmids encoding the indicated proteins were lysed in IAA (25 mM) and visualized by immunoblotting. In **B**-**E**, reducing refers to the presence of DTT (50 mM) in the loading dye for SDS-PAGE analysis.

We initially transiently transfected HEK 293T cells with a construct encoding wild-type NLRP1-FLAG or NACHT-LRR-FLAG before lysing in either IAA or NEM, performing anti-FLAG IP, and visualizing complexes by non-reducing SDS-PAGE analysis (**Fig. 3B**,**C**). Notably, no NLRP1-TRX1 conjugate was observed in the absence of IAA or NEM, showing that the two proteins do not appear to form a stable linkage natively. Intriguingly, however, these experiments revealed the presence of new bands that migrated slightly slower than the corresponding FLAG-tagged protein in the presence of NEM and IAA (**Fig. 3B**,**C**, bottom gels). Consistent with stable association with TRX1, these higher molecular weight bands were also reactive toward a TRX1 antibody (**Fig. 3B**, top gel). It should be emphasized that not all full-length (FL) NLRP1 undergoes autoproteolysis, so the TRX1 antibody detects two bands that correspond to the TRX-NLRP1^FL^ and the TRX-NLRP1^NT^ conjugates. Moreover, disulfide bonds clearly mediated these intermolecular interactions, as H_2_O_2_ increased and DTT abrogated their appearance (**Fig. 3B**,**C**).

Overall, these data indicate that NLRP1 and TRX1 can form a transient disulfide bond. Although both IAA and NEM can trap this conjugate, IAA is likely more effective due to its smaller size.

To confirm that C427^NLRP1^/C225 ^NLRP1B^ mediated the intermolecular disulfide bond between TRX1-NLRP1, we performed similar thiol trapping experiments using various NLRP1 cysteine mutants. (**Fig. 3D**,**E**). As expected, only human NLRP1^C427S^ and mouse NLRP1B^C225S^ did not form the TRX1-NLRP1 conjugate (**Fig. 3D**,**E**). In addition, we previously showed that mutation of the Walker B motif (D413A) in the ATPase active-site of the NACHT domain abrogates TRX1 binding, possibly by destabilizing the folding of the domain ^13^. Consistent with these data, we found that NLRP1^D314A^ did not form a stable TRX1 conjugate in a trapping experiment. We should note that TRX1^C35S^, which weakly binds to NLRP1^13^, similarly forms a conjugate with NLRP1^WT^, but not NLRP1^C427S^, on non-reducing gels. We reason that the lack of C35 in the TRX1^C35S^-NLRP1^WT^ conjugate, like the IAA- and NEM-trapped conjugates, prevents its resolution into two separate species once formed (**Fig. S3A**).

Intriguingly, C427 of NLRP1 aligns with C319 of NLRP3 (**Fig. S3B**). Although we previously showed that NLRP3 does not associate with TRX1^13^, we wondered if a transient or weaker affinity interaction between TRX1 and NLRP3 might be revealed using the IAA-trapping protocol (**Fig. S3C**). However, we found that NLRP3, unlike NLRP1, does not form intermolecular with TRX1 even after IAA treatment. Thus, NLRP3 does not associate with TRX1.

### Mass spectrometry identification of the adduct

Although these experiments show that C427^NLRP1^ forms a disulfide bond with TRX1, we nevertheless wanted to confirm these findings using LC-MS/MS (**Fig. 4A**). To accomplish this, we transiently transfected HEK 293T cells with constructs encoding HA-tagged TRX1 and FLAG-tagged NACHT-LRR-FLAG, lysed cells in the presence of NEM, and enriched proteins by anti-FLAG IP. As expected, Coomassie staining of the SDS-PAGE gel revealed two bands at ∼75 and ∼100 kDa corresponding to the NACHT-LRR and NACHT-LRR-TRX1 conjugate, respectively.

**Figure 4.**
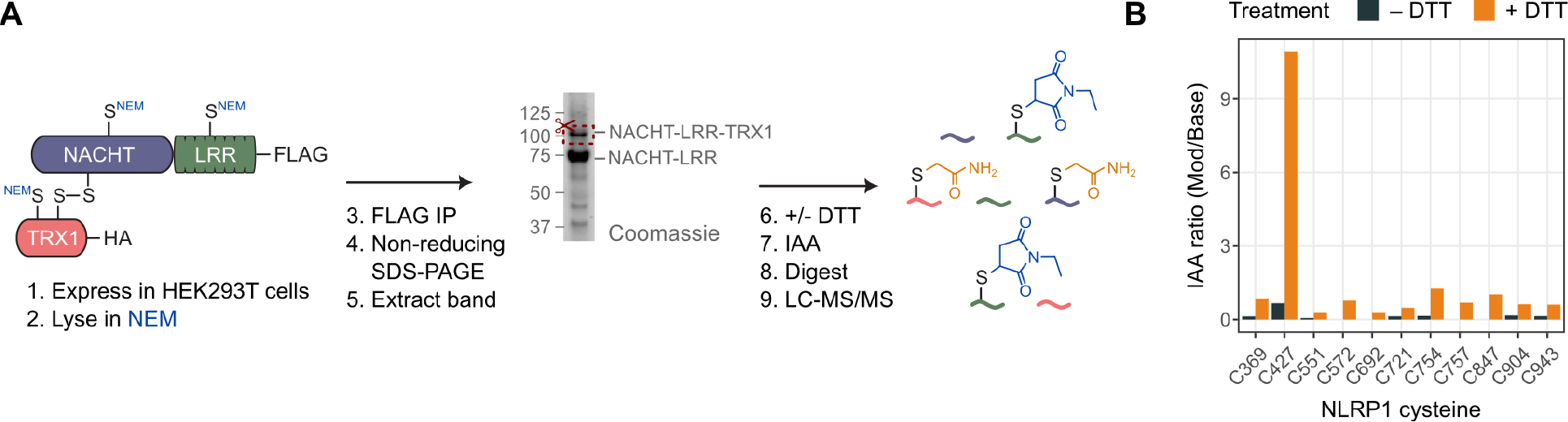
Mass spectrometry-based identification of the modified cysteine residue. (**A**) Schematic of the workflow used to identify cysteine residues involved in the NLRP1-TRX1 disulfide bond by LC-MS/MS. The excised gel band is shown. (**B**) Ratio of extracted ion current (XIC) for peptides with IAA-modified cysteine residues compared to the total XIC for the corresponding identified peptide sequence, following the schematic in **A**.

The band corresponding to the conjugate was excised, reduced with DTT, labeled with IAA, enzymatically digested with trypsin, Glu-C, and Asp-N, and subjected to LC-MS/MS analysis. This workflow reduces and caps any disulfide bonds in the excised band with IAA, thereby making them distinct from the reduced sulfhydryl groups that were originally alkylated by NEM. This experiment revealed that only NLRP1 C427 was highly modified by IAA after DTT reduction (**Fig. 4B**). Overall, these data further confirm that C427^NLRP1^ forms a transient disulfide with C32^TRX1^ that can be trapped with thiol-reactive agents.

## Discussion

We first reported that the oxidized form of TRX1 binds to and represses the NLRP1 inflammasome more than two years ago ^12,13^. However, it remained unknown how NLRP1 selectively associates with only the oxidized, but not the reduced, form of TRX1, which, apart from the disulfide bond, have highly similar structures ^19^. Here, we leveraged AlphaFold-Multimer to predict the structure of the TRX1-NLRP1 complex, which strikingly suggested that a specific cysteine in both human and mouse NLRP1 NACHT domains associates with the TRX1 active site. Using site-directed mutagenesis, thiol-trapping experiments, and mass spectrometry, we revealed that this cysteine on NLRP1 forms a transient disulfide bond with oxidized TRX1, thereby acting as a selectivity filter for the oxidized form of TRX1.

During the preparation of this manuscript, a publication appeared in *Nature* reporting the cryo-EM structure of the human NLRP1 NACHT-LRR region in complex with TRX1 from *Spodoptera fugiperda* ^20^. Notably, the AlphaFold-Multimer prediction described here is highly similar to this solved cryo-EM structure, showcasing the power of AlphaFold-Multimer to illuminate protein-protein interfaces, including those with unusual disulfide bonds. In addition to reporting this new structure, this recent manuscript largely confirmed our prior work that oxidized TRX1 binds to and represses the activation of the NLRP1 inflammasome ^12,13^.

Projecting forward, many mysteries remain regarding the NLRP1-TRX1 interaction. First, it is not precisely clear how TRX1 stabilizes the folding of the NACHT-LRR. The AlphaFold prediction of the isolated NACHT-LRR domain suggests that an ordered loop (“loop 1”, residues 288-311) in the NLRP1-TRX1 structure is disordered in the TRX1-unbound form (**Fig. S4**). Future studies are needed to determine if the unraveling of this loop, or potentially a different “loop 2” (**Fig. S4**), accelerates NT degradation. Second, the key proteins that recognize and degrade NLRP1 have not yet been identified. We speculate that TRX1 binding may slow degradation by the core 20S proteasome, which degrades misfolded proteins independent of ubiquitin^21-23^, but additional studies are needed to confirm that this pathway is operational. Lastly, and most importantly, it is not yet clear why an oxidized cell state represses NLRP1 inflammasome activation. We hypothesize two general possibilities: 1) that pathogens induce some form of reductive stress that accelerates NT degradation, or 2) that oxidized TRX1 serves to restrict inflammasome activation to a reduced metabolic state. Considerably more effort is needed to resolve these outstanding points and fully characterize the complex relationship between the intracellular redox state and the innate immune system.

## Acknowledgements

This work was supported by the NIH (R01 AI137168, R01 AI163170, R01 CA266478 to D.A.B.; the MSKCC Core Grant P30 CA008748), Mr. William H. and Mrs. Alice Goodwin, the Commonwealth Foundation for Cancer Research, and The Center for Experimental Therapeutics of Memorial Sloan Kettering Cancer Center (D.A.B.), the Technology Development Fund of Memorial Sloan Kettering Cancer Center (D.A.B.), and The Starr Foundation Basic Research Innovation Award Grant (M.B.G). We thank Juliana Ortiz-Pacheco and Mara Monetti of the MSK Proteomics Core for discussions and assistance with mass spectrometry-based assays.

## Author Contributions

M.B.G., J.C.H., L.P.T., and D.P.B. performed cloning, gene editing, biochemistry, and cell biology experiments. All authors designed experiments and analyzed data. D.A.B. directed the project. D.A.B., M.B.G., and J.C.H. wrote the manuscript.

## Materials and Methods

### Antibodies and reagents

Antibodies used include: Thioredoxin 1 (ab86255, Abcam; 2429S, Cell Signaling Technology), DPP9 (ab42080, Abcam), GAPDH (2118S, Cell Signaling Technology), HA rabbit monoclonal Ab (Cell Signaling Tech, C29F4), HA mouse monoclonal Ab (Cell Signaling Tech, 6E2), FLAG mouse monoclonal Ab (Sigma Aldrich, F1804), IRDye 800CW donkey anti-rabbit (LICOR, 925-32211), IRDye 680RD donkey anti-rabbit (925-68073), IRDye 800CW donkey anti-mouse (925-32212), IRDye 680RD donkey anti-mouse (925-68072). Reagents for immunoprecipitation experiments include anti-FLAG M2 agarose resin (Millipore, A2220). Other reagents used include doxycycline hyclate (Cayman, 14422), FuGENE HD (Promega, E2311), HALT protease inhibitor cocktail (Thermo Scientific, 78430), Iodoacetamide (Thermo Scientific, A39271), N-ethylmaleimide (Thermo Scientific, 23030).

### Cell culture

HEK 293T cells were grown in Dulbecco’s Modified Eagle’s Medium (DMEM) with L-glutamine and 10% fetal bovine serum (FBS). THP-1 cells were grown in Roswell Park Memorial Institute (RPMI) medium 1640 with L-glutamine and 10% FBS. All cells were grown at 37 °C in a 5% CO_2_ atmosphere incubator. Cell lines were regularly tested for mycoplasma using the MycoAlert Mycoplasma Detection Kit (Lonza). *CARD8*^*-/-*^THP-1 cells were generated as previously described ^13,24,25^. Doxycycline (DOX)-inducible NLRP1 WT and mutant knock-in *CARD8*^*-/-*^ THP-1 were generated as previously described ^9,26^.

### Transient transfections

HEK 293T cells were plated in 6-well tissue culture plates at 5.0 × 10^5^ cells/well in DMEM. The next day, the indicated plasmids were mixed with an empty vector to a total of 2.0 μg DNA in 125 μL Opti-MEM and transfected using FuGENE HD (Promega) according to the manufacturer’s protocol. For larger scale experiments, the above protocol was repeated with HEK293T cells at 3.0× 10^6^ cells/10 cm plate and the transfection performed with 20.0 μg DNA in 330 μL Opti-MEM. For experiments involving constitutive plasmid expressions, cells were incubated for an additional 48 hours before their harvest. For experiments involving tet-inducible plasmid constructs, cells were treated with DOX at 1 ug/mL (and/or with other compounds) 20 hours after transfection then incubated for an additional 24 hours before harvesting.

### Cloning

Plasmids for human NLRP1 WT, mouse NLRP1B WT, and their variants were cloned as described previously ^9,24,26^. Briefly, DNA sequences encoding the genes were purchased from GenScript, amplified by polymerase chain reaction (PCR), shuttled into the Gateway cloning system (ThermoFisher Scientific) using pDONR221, pLEX307, and pInducer20 vectors from Addgene. Site-directed mutagenesis was performed using the QuikChange II Site-directed mutagenesis kit (Agilent) according to the manufacturer’s instructions.

### Immunoprecipitation and immunoblotting

Cell pellets obtained from transfected HEK 293T cells were sonicated (5 seconds, 10% amplitude) in PBS. For conjugate trapping experiments, iodoacetamide (IAA) or N-ethylmaleimide (NEM) were added at the indicated concentrations into PBS before sonication. The samples were then incubated in the dark for 30 minutes shaking at room temperature. The clarified lysates were retained (centrifuged at 1000 x *g* for 5 min), normalized using the DCA Protein Assay Kit (Bio-Rad), and were subsequently incubated with 40 μL of anti-FLAG-M2 agarose resin (Sigma) per sample for 1 hour at room temperature. After washing with two rounds of 100 uL of PBS, bound proteins were eluted by incubating resin with 100 μL of PBS containing 150 ng/μL 3X-FLAG peptide for 1 hour at room temperature. For subsequent immunoblotting assays, the eluates were added with loading dye containing either 50 mM of dithiothreitol (DTT) for reducing conditions or none for non-reducing conditions. Samples were then incubated at 95 °C for 10 min. The samples were separated by SDS-PAGE, immunoblotted, and visualized using the Odyssey Imaging System (Li-Cor).

### Gel Extraction of the NLRP1-TRX1 conjugate

HEK 293T cells pellets transfected with plasmids encoding NACHT-LRR-FLAG and TRX1-HA were sonicated in PBS containing 25 mM NEM. Proteins were enriched using FLAG agarose beads and separated by SDS-PAGE in non-reducing conditions. The membrane was then stained with SimplyBlue SafeStain (Invitrogen) and visualized under light. Replicates of NACHT-LRR-TRX1 conjugate bands were then excised and digested following a standard in-gel digestion protocol. During the reduction and alkylation steps, only one group received 10 mM DTT dissolved in 25 mM NH_4_HCO_3_, while the other received only NH_4_HCO_3_. Then, both groups were treated with 50 mM iodoacetamide. The gel pieces were subsequently proteolytically digested overnight with trypsin, Glu-C, and Asp-N at 37 °C. The peptides were extracted and desalted with C18 Disk Filters (3M Empore).

### Mass spectrometry and analysis

Peptides were separated on an ACQUITY Peptide BEH C18 Column (130 Å, 1.7 um, 75um × 250mm) coupled to an ACQUITY UPLC M-Class C18 Trap Column (100 Å, 5um, 2G,V/M, 180um × 200mm) using a gradient from 1% to 30% B over 110 minutes and then 50 to 90% B for 15 minutes (Buffer A: 0.1% Formic Acid in HPLC grade water; Buffer B: 99.9% Acetonitrile/0.1% Formic Acid) with a flow rate of 300 nL/min using a NanoAcquity system (Waters). MS data were acquired on a QExactive Plus mass spectrometer (Thermo Scientific) using data-dependent acquisition top 10 method, ACG target of 1e6, maximum injection time of 50 msec, scan range 380 to 1600 m/z and a resolution of 70K. MS/MS was performed at a 17.5K resolution, ACG target 5e4, maximum injection time of 50 msec, isolation window of 1.5 m/z. Dynamic exclusion was set at 15 sec. Thermo RAW files were analyzed using MAXQuant (v 2.1.0). Oxidation (M), deamidation (NQ), IAA (C), N-ethyl maleimide (C) and hydrolyzed N-ethyl maleimide (C) were included as variable modifications.

### Immunoblotting

Cells were washed 2 × in PBS (pH = 7.4), resuspended in PBS, and lysed by sonication. Protein concentrations were determined and normalized using the DCA Protein Assay kit (Bio-Rad). Clarified lysates (5 min × 5,000 *g*) were then suspended in 1X PBS containing HALT protease inhibitor (Thermo) and combined 1:1 with 2X sample loading buffer (DTT added as indicated) before heating to 95 °C for 10 min. Samples were run on NuPAGE 4 to 12%, Bis-Tris, 1.0 mM, Midi Protein Gel (Invitrogen) for 45 - 60 min at 175 V. Gels were transferred to nitrocellulose with the Trans-Blot Turbo Transfer System (BIO-RAD). Membranes were blocked with Intercept (TBS) Blocking Buffer (LI-COR) for 30 min at ambient temperature, before incubating with primary antibody overnight at 4°C. Blots were washed 3 times with TBST buffer before incubating with secondary antibody for 60 min at ambient temperature. Blots were washed 3 times, rinsed with water, and imaged via Odyssey CLx (LI-COR).

### AlphaFold-Multimer Predictions

AlphaFold-Multimer was accessed via the Cosmic2 portal (https://cosmic-cryoem.org). FASTA files were generated for the human NLRP1 NACHT-LRR domains (residues K229 to S990; UniProt ID: Q9C000) or mouse NLRP1B (allele 1, residues K30 to V810; Uniprot ID Q2LKW6). The sequence for human TRX1 (Uniprot ID P10599) was used in both models. Alphafold-Multimer models were generated using the default options and the resulting .pdb files were inspected using Pymol.

## Supplementary Figures

**Figure S1.**
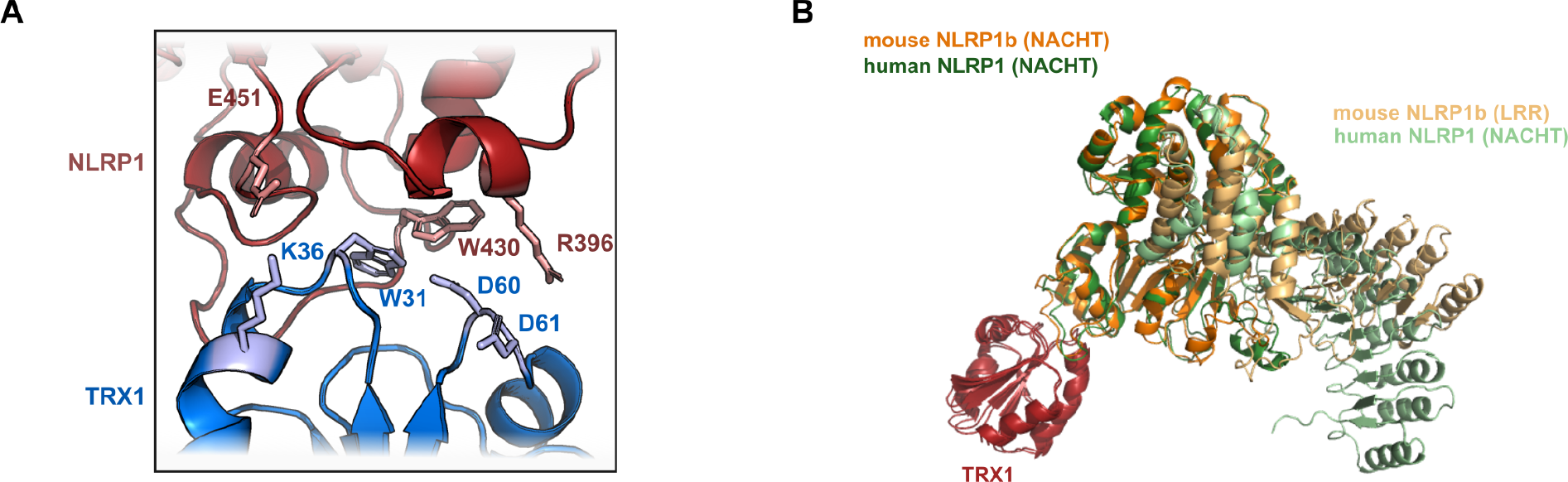
Additional views of the predicted NLRP1-TRX1 interaction. (**A**) Selected contacts and residue side chains at the AlphaFold-Multimer predicted TRX1-NLRP1 interface. (**B**) Overlay of the AlphaFold-Multimer predicted models for TRX1-NLRP1 (human) and TRX1-NLRP1B (mouse).

**Figure S2.**
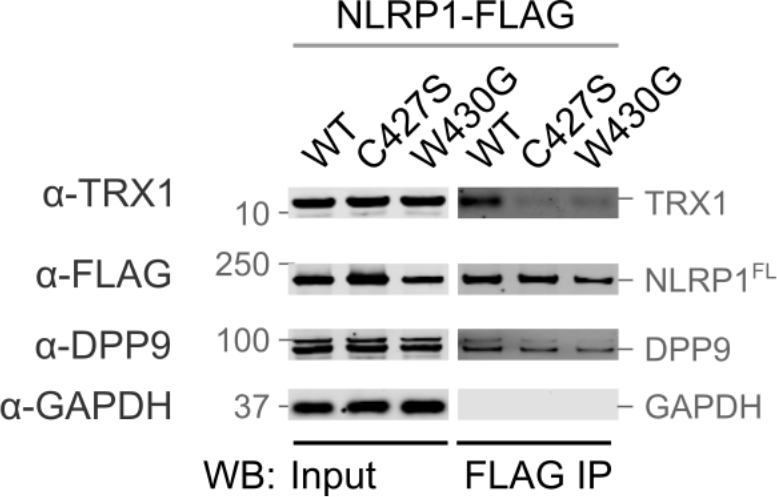
W430 within the NLRP1 hydrophobic ridge is critical for TRX1 binding. HEK 293T cells were transiently transfected with plasmids encoding NLRP1-FLAG constructs before lysates were subjected to anti-FLAG IP and immunoblotting analyses.

**Figure S3.**
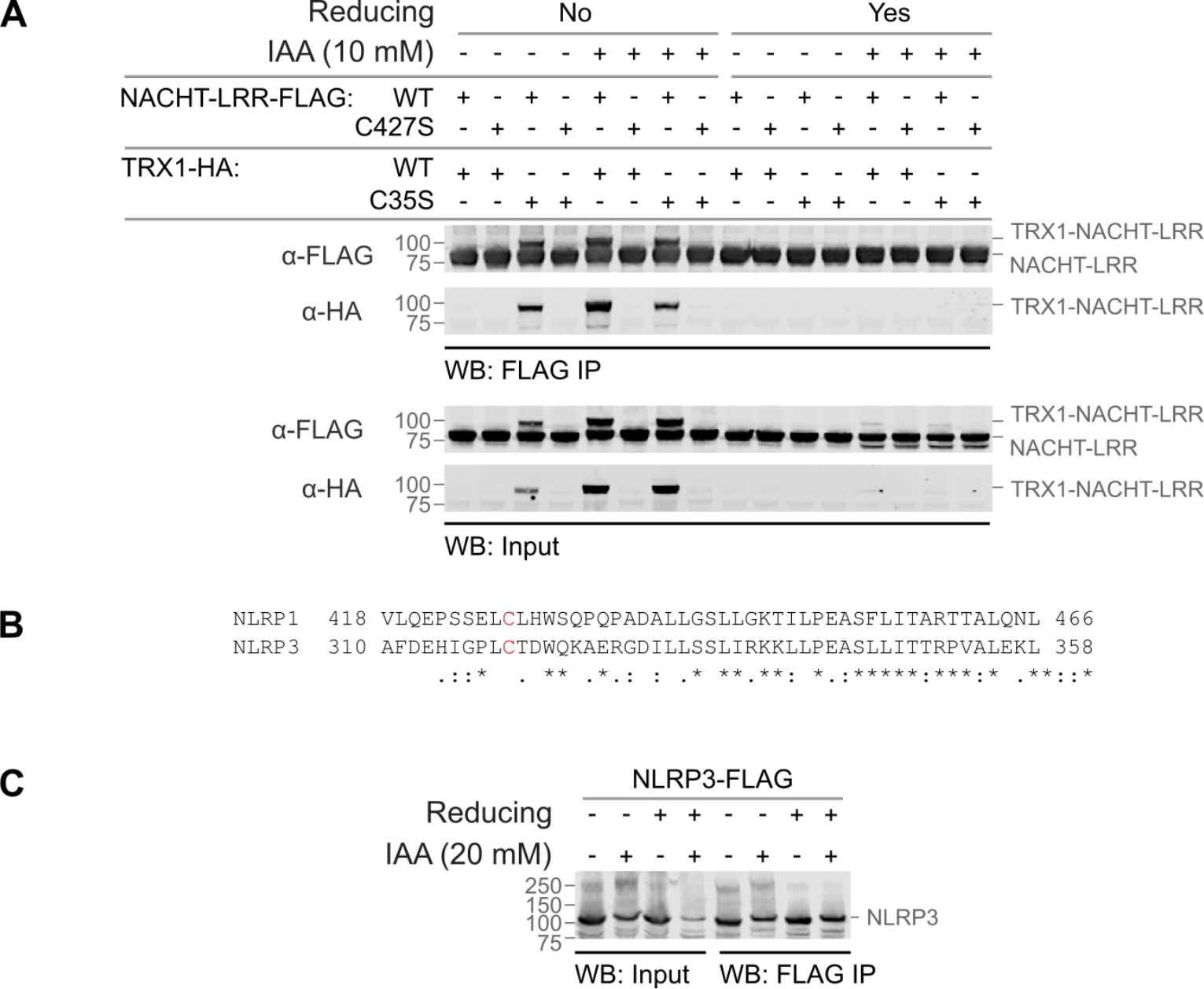
C35S mutant TRX1 forms a conjugate with C427 of NLRP1. (**A**) HEK 293T cells were transiently transfected with plasmids encoding the indicated NACHT-LRR-FLAG and TRX1-HA constructs and lysed in IAA. Proteins were then enriched by anti-FLAG IP and analyzed by immunoblotting. (**B**) Sequence alignment of NLRP1 and NLRP3 around the conserved cysteine (C427 in NLRP1), which is colored red. (**C**) HEK 293T cells were transiently transfected with a plasmid encoding NLRP3-FLAG before lysates were subjected to anti-FLAG IP and immunoblotting analyses.

**Figure S4.**
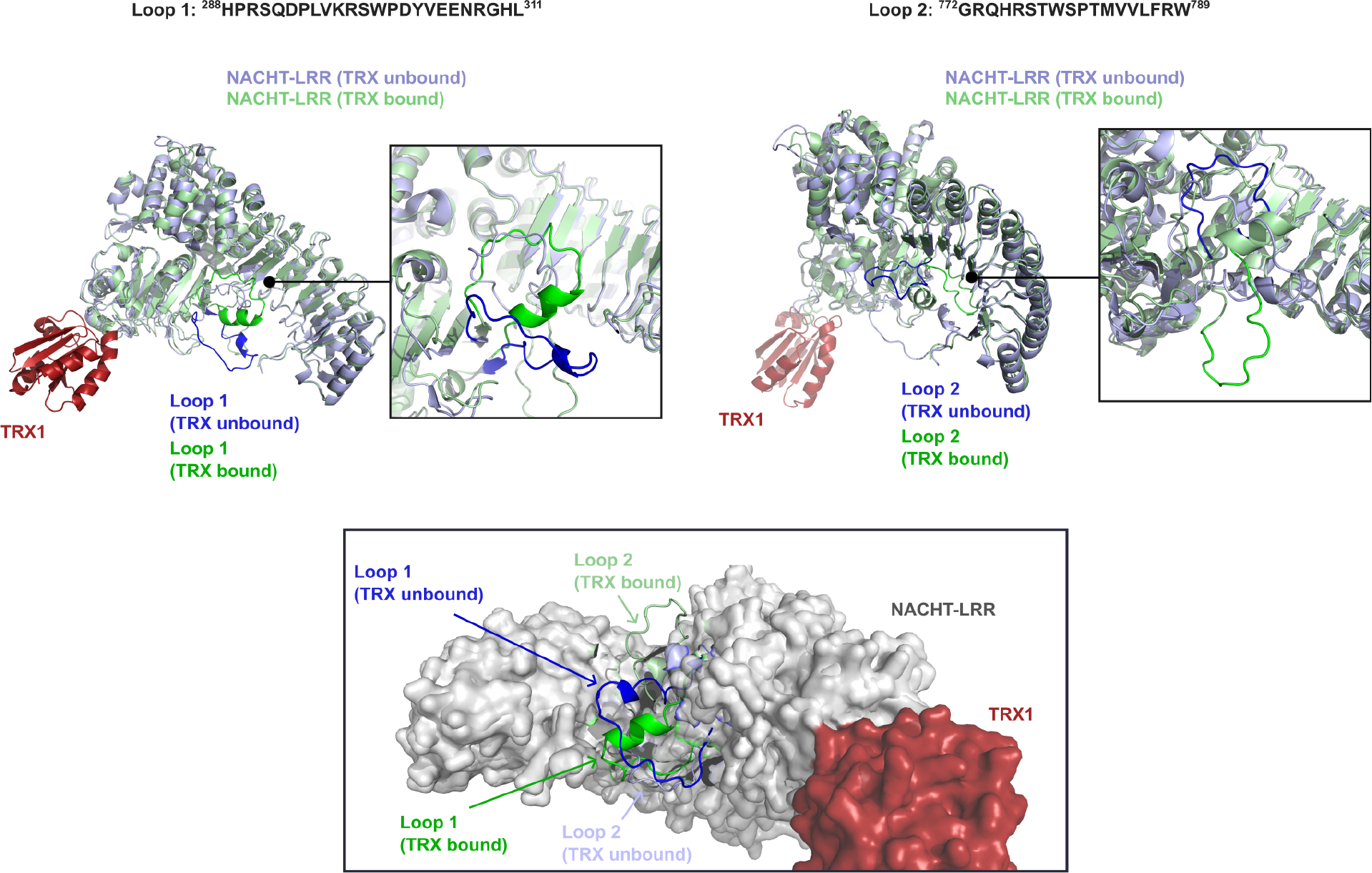
AlphaFold-Multimer predicts that TRX1 orders at least one loop within the NLRP1 NACHT-LRR. The AlphaFold prediction for the unbound NLRP1 NACHT-LRR was aligned with the AlphaFold-Multimer model of the TRX1-NLRP1 complex described in **Figure 1**. This analysis suggests that two loops (Loop 1 and Loop 2) potentially undergo dramatic conformational changes upon TRX1 binding.

